# PGC1/PPAR Drive Cardiomyocyte Maturation through Regulation of Yap1 and SF3B2

**DOI:** 10.1101/2020.02.06.937797

**Authors:** Sean Murphy, Matthew Miyamoto, Anais Kervadec, Suraj Kannan, Emmanouil Tampakakis, Sandeep Kambhampati, Brian Leei Lin, Sam Paek, Peter Andersen, Dong-Ik Lee, Renjun Zhu, Steven S. An, David A. Kass, Hideki Uosaki, Alexandre R. Colas, Chulan Kwon

## Abstract

Cardiomyocytes undergo significant levels of structural and functional changes after birth—fundamental processes essential for the heart to produce the volume and contractility to pump blood to the growing body. However, due to the challenges in isolating single postnatal/adult myocytes, how individual newborn cardiomyocytes acquire multiple aspects of mature phenotypes remains poorly understood. Here we implemented large-particle sorting and analyzed single myocytes from neonatal to adult hearts. Early myocytes exhibited a wide-ranging transcriptomic and size heterogeneity, maintained until adulthood with a continuous transcriptomic shift. Gene regulatory network analysis followed by mosaic gene deletion revealed that peroxisome proliferator-activated receptor coactivator-1 signaling—activated in vivo but inactive in pluripotent stem cell-derived cardiomyocytes—mediates the shift. The signaling regulated key aspects of cardiomyocyte maturation simultaneously through previously unrecognized regulators, including Yap1 and SF3B2. Our study provides a single-cell roadmap of heterogeneous transitions coupled to cellular features and unveils a multifaceted regulator controlling cardiomyocyte maturation.

**Significance Statement:** How the individual single myocytes achieve full maturity remains a ‘black box’, largely due to the challenges with the isolation of single mature myocytes. Understanding this process is particularly important as the immaturity and early developmental arrest of pluripotent stem cell-derived myocytes has emerged a major concern in the field. Here we present the first study of high-quality single-cell transcriptomic analysis of cardiac muscle cells from neonatal to adult hearts. We identify a central transcription factor and its novel targets that control key aspects of myocyte maturation, including cellular hypertrophy, contractility, and mitochondrial activity.

---

Decades of advances in cellular and developmental cardiology have provided fundamental insights into understanding myocardial lineage specification in vivo, and this knowledge has been instrumental for producing cardiomyocytes (CMs) from pluripotent stem cells (PSCs) (Devalla and Passier, 2018; Evans et al., 2010; Kattman et al., 2011). However, while newborn CMs continue to increase their volume and contractility through extensive morphological, functional, and metabolic changes until adulthood, PSC-derived CMs (PSC-CMs) are mired in an immature state even after long-term culture (DeLaughter et al., 2016; Kannan and Kwon, 2018; Uosaki et al., 2015). The lack of maturity significantly limits scientific and therapeutic applications of PSC-CMs. Furthermore, despite a number of genes involved in CM maturation are associated with cardiomyopathies, little is known about its relevance to the initiation and progression of cardiac pathogenesis (Shenje et al., 2014; Uosaki et al., 2015). Thus, there is a significant need to understand biological processes underlying CM maturation in vivo.

CM maturation is a complex process essential for the heart to circulate blood to the rapidly growing body (Yang et al., 2014). After terminal differentiation, CMs undergo binucleation/polyploidization around the first week of birth in mice. They gradually increase in size and become rectangular with uniformly patterned sarcomeres (Hirschy et al., 2006). To efficiently propagate electrical activity, the plasma membrane invaginates into the cells and forms transverse tubules, enabling excitation-contraction coupling (Ziman et al., 2010). The myocytes become tightly connected via intercalated discs to allow simultaneous contraction. These events are accompanied with functional and metabolic changes including mature calcium handling, increased contractile force, and mitochondrial maturation and oxidative phosphorylation (Yang et al., 2014). These multi-adaptive changes occur in the early postnatal period and continue until adolescent/adult stages. However, it remains an open question whether these distinct processes occur in a coordinated fashion. Factors and pathways mediating these individual processes are poorly understood as well.

Previous large-scale meta-analyses provided a transcriptomic atlas of cardiac maturation, allowing us to determine gene regulatory networks and pathways involved in cardiac maturation (Uosaki et al., 2015). The scope was, however, largely focused on prenatal stages, leaving the postnatal transcriptome dynamics unclear. Moreover, cell-to-cell variations—poorly understood in the field—could not be determined with bulk analysis. This issue can be addressed by single-cell RNA-sequencing (scRNA-seq) that enables comprehensive analysis of developmental and cellular trajectory and heterogeneity (Grun and van Oudenaarden, 2015). However, scRNA-seq is rarely utilized in myocyte biology due to technical difficulties associated with single-cell isolation of healthy, mature CMs.

We have recently demonstrated that large-particle fluorescence-activated cell sorting (LP-FACS) enables high-quality scRNA-seq and functional analysis of mature adult CMs (Kannan et al., 2019). Based on this, here we present high-quality scRNA-seq analysis of CMs isolated from neonatal to adult hearts. We demonstrate that newborn CMs are highly heterogeneous and progressively change expression of genes regulating cellular hypertrophy, contractility, and metabolism until adulthood. By combining gene regulatory network analysis with mosaic gene deletion and chromatin immunoprecipitation sequencing (ChIP-seq), we identify peroxisome proliferator-activated receptor (PPAR) coactivator-1 (PGC1) signaling as a multi-faceted regulator coordinating CM maturation via its novel targets Yap1 and SF3B2.

## Results

### CMs Exhibit High Levels of Transcriptomic Heterogeneity During Postnatal Maturation

Despite serving as a powerful tool in biology and medicine, the large size and fragility of mature CMs have limited the use of scRNA-seq in studying CM growth and disease (Ackers-Johnson et al., 2018). Conventional sorting or microfluidic platforms result in damaged/ruptured myocytes due to inappropriate nozzle size/ flow rate, leading to abnormal transcript reads and cell death. To address this, we recently tested LP-FACS and found that it allows isolation of healthy, mature myocytes, enabling both high-quality scRNA-seq and functional analysis (Kannan et al., 2019). Utilizing LP-FACS, we asked how postnatal CMs become mature cells at the single cell transcriptome level. We harvested hearts from postnatal day (p) 0, p7, p14, p21, and p28 mice and dissociated CMs using standard Langendorff perfusion, followed by isolation of viable single myocytes with LP-FACS (Kannan et al., 2019) (Figure 1A). Curiously, tSNE (van der Maaten and Hinton, 2008)-based clustering of p0–p28 single cell samples show partial segregation (Figure 1B), suggesting that significant numbers of cells at each stage may have similar transcriptome profiles as those of cells present at the other stages. We then used Monocle2 (Qiu et al., 2017) to organize transcriptome profiles of CMs at different developmental stages based on transcriptomic similarities. The Monocle-based analysis produced a single pseudotime trajectory showing a progressive, unidirectional pattern of CMs over the course of maturation (Figure 1C). Strikingly, individual cells from the same stages were found distributed broadly over the trajectory (Figure 1D). When comparing the maturation score (as indicated by pseudotime) among different timepoints, we see that while average maturation scores increase with age, transcriptomic heterogeneity is maintained even at p28 myocytes (Figure 1D). In agreement with this, similar levels of cell size heterogeneity were found in postnatal CMs (Figure 1E).

**Fig. 1.**
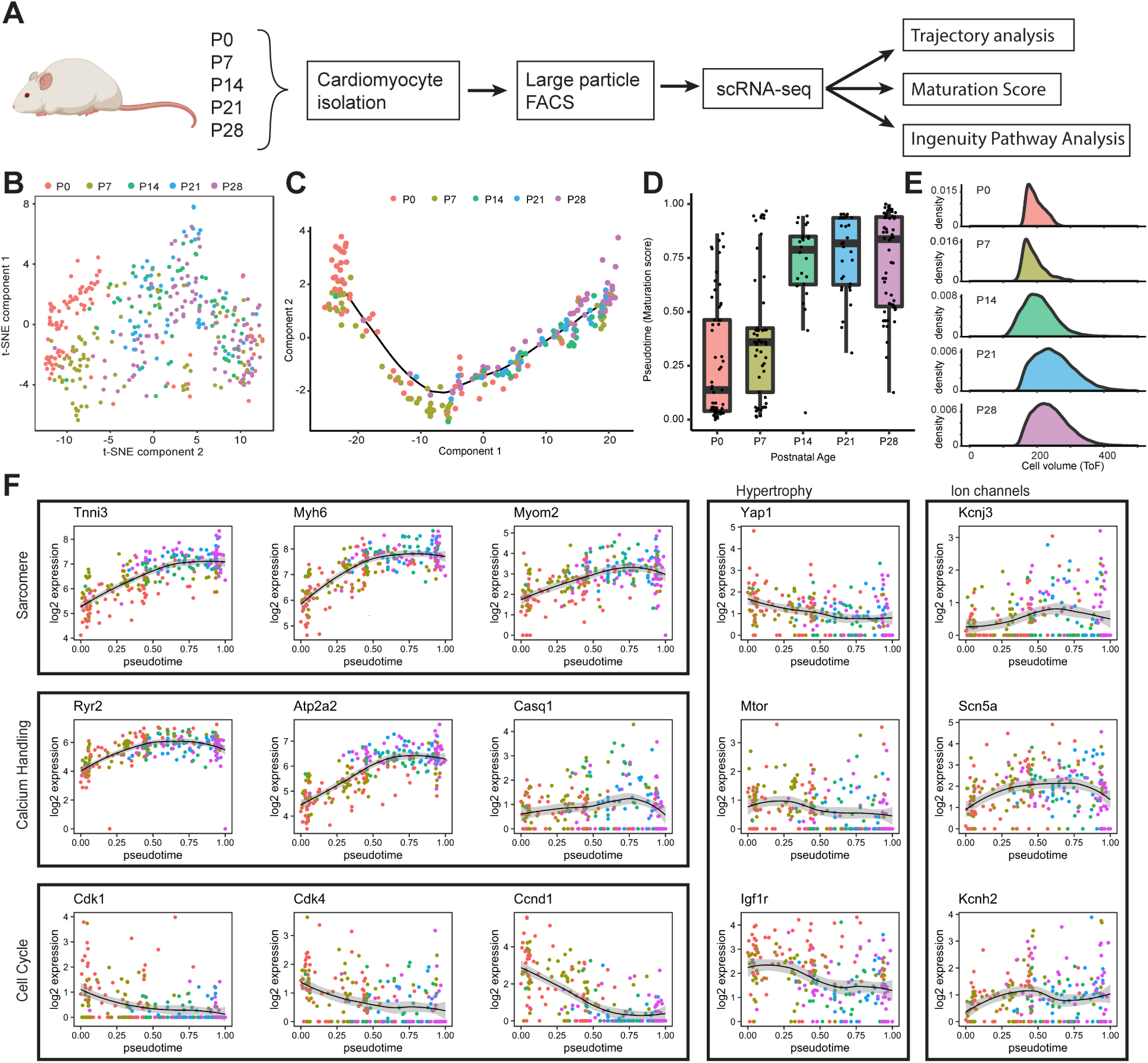
Postnatal CMs exhibit high levels of transcriptomic heterogeneity. A, Experimental design for scRNA-seq and computational analysis of CMs isolated from p0–p28 hearts. B, t-SNE plot representation of p0–p28 CMs. C, Monocle-based developmental trajectory of p0–28 CMs. D, Distribution of normalized pseudotimes (maturation scores) by age. E, FACS-based cell size analysis with time of flight. F, Log expression of CM genes associated with structural maturation, calcium handling, cell cycle, hypertrophy, and ion channels plotted over pseudotime.

We next examined the expression profiles of maturation-associated genes by plotting them along pseudotime, grouped by function. Expression levels of structural genes known to be upregulated in mature CMs, including Myh6, Tnni3, Myom2 (Yang et al., 2014), were gradually increased (Figure 1F). Calcium handling genes and ion channels including Ryr2, Atp2a2, Casq1, critical for contractility development, were continually upregulated while genes involved in cell cycle, including Cdk1, Cdk4, Ccnd1 (Mohamed et al., 2018; Yang et al., 2014), were downregulated (Figure 1F). Conserved genes governing cellular hypertrophy, including Mtor, Yap1, Igf1r (Lloyd, 2013; Perez-Gonzalez et al., 2019), were modestly downregulated or maintained in expression as they are known to regulate cell volume (Figure 1F). Gene ontology (GO) analysis indicated that genes related to muscle contraction and cellular metabolism are highly regulated during the process (Figure S1A). Consistently, hierarchical clustering showed that mitochondrial gene expression is gradually increased over time (Figure S1B). Our single-cell approach quantitatively shows that postnatal CM maturation takes place in a continuous, but highly heterogeneous fashion.

### Gene Network Analysis Predicts PGC1/PPAR Signaling as A Key Regulator of Cardiac Maturation

The regulatory mechanisms underlying cardiomyocyte maturation are largely unknown. To gain mechanistic insights into upstream regulators governing postnatal CM maturation, we used Ingenuity Pathway Analysis that infers regulators of differentially expressed genes by knowledge base of expected effects between transcriptional regulators and their target genes (Kramer et al., 2014). Using the scRNA-seq dataset, we first generated gene regulatory networks with predicted upstream regulators by p-value and activation scores. We found that transcriptional cofactors and a group of nuclear receptors (PPARs, thyroid hormone receptors, retinoid receptors, etc) are predicted to be upstream transcriptional regulators significantly affecting the overall gene expression changes during maturation (Figure 2A, Figure S1C, Figure S1D, and Figure S1E). We further used a cut off of 10e-5 for the false discovery rate and ranked them by p-value. This analysis inferred the transcriptional cofactors PGC1*α*/*β*—two PGC1 isoforms—as among the most influential factors (Figure 2C). Thus, we quantified their expression levels in developing hearts and PSC-CMs. Expression of the cofactors and nuclear receptors was gradually increased from embryonic to adult stages in vivo, with more pronounced upregulation after birth (Figure 2B, d). An incremental expression pattern was also observed in most of the genes in long-term cultured PSC-CMs, but PGC1/PPAR*α* levels remained constantly low (Figure 2B). This indicates that PGC1/PPAR*α* are misregulated in PSC-CMs and may be responsible for their maturation arrest (DeLaughter et al., 2016; Uosaki et al., 2015).

**Fig. 2.**
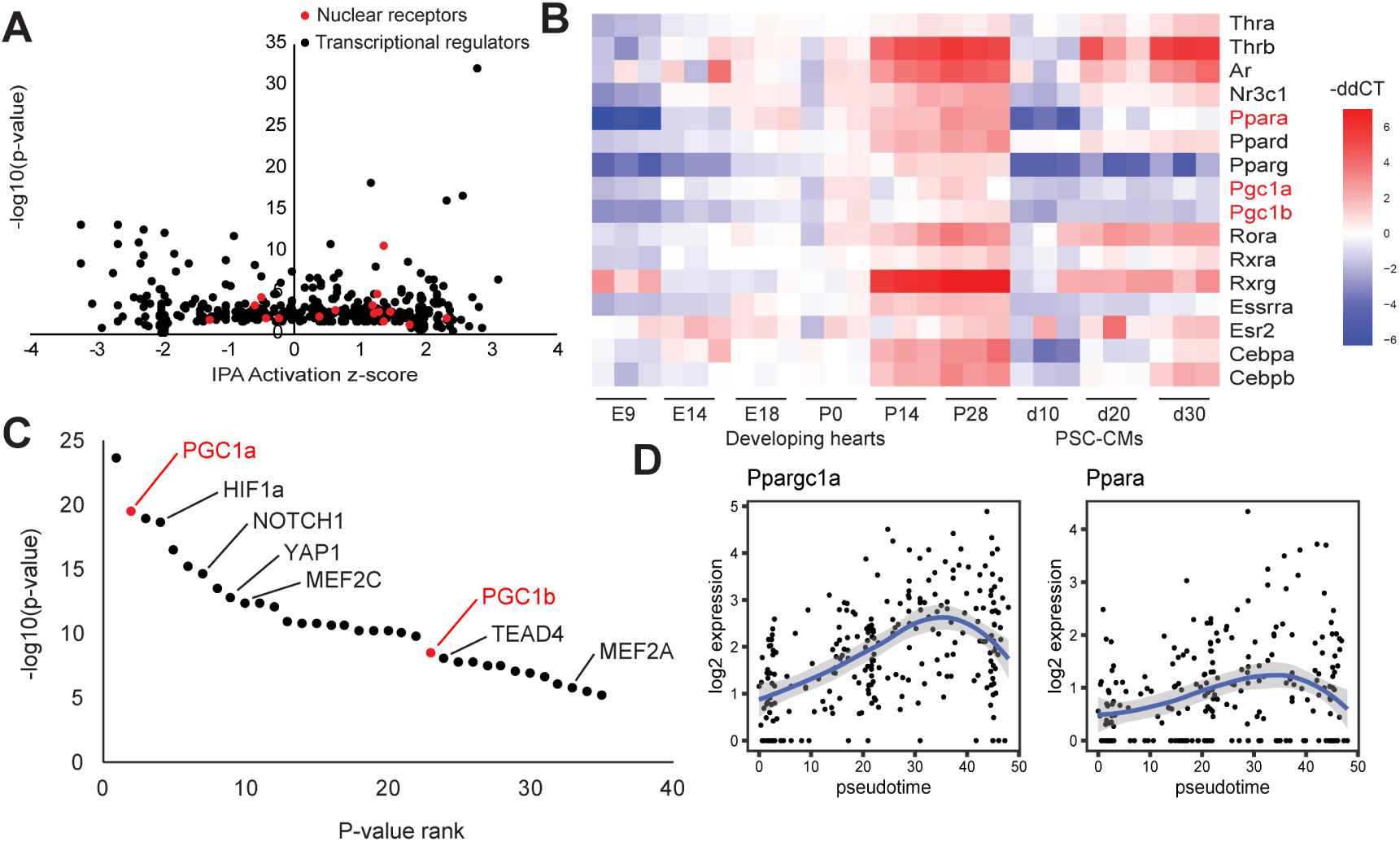
PGC1/PPAR is a predicted key upstream regulator of CM maturation. A, Top transcriptional regulators plotted by p-value and IPA activation z-score with nuclear receptors highlighted in red. B, Heatmap of gene expression of PGC1 and nuclear receptors in developing mouse hearts and cultured PSC-CMs, quantified by qPCR. C, P-value ranking of top upstream regulators of CM maturation with two PGC1 isoforms highlighted in red. D, Expression trends over pseudotime of PGC1 and PPAR*α* in postnatal CMs.

### PGC1 Is Required Cell-Autonomously for Postnatal CM Growth and Contractility Development

PGC1*α/β* are conserved transcriptional coactivators for PPARs and other nuclear receptors and known as central regulators of energy metabolism (Finck and Kelly, 2006). The two isoforms show extensive sequence homology with functional redundancy (Lai et al., 2008; Rowe et al., 2010). In fact, embryonic deletion of either PGC1*α* or PGC1*β* does not affect heart formation, but the double knockout results in lethality soon after birth with small hearts accompanied by mitochondrial defects (Lai et al., 2008). However, these studies deleted the alleles globally or in embryonic stages, leaving their cell-autonomous, postnatal role unknown. Based on the single cell bioinformatics prediction and low levels of expression in PSC-CMs, we hypothesized that PGC1 intrinsically mediates postnatal maturation of CMs. To test this, we generated conditional mosaic knockout (cmKO) mice, which avoids non-cell-autonomous effects and early lethality caused by global or conditional deletion (Figure 3A). For this, we generated PGC1/alpha//beta flox/flox; Ai9 mice and administered AAV vectors expressing Cre specifically in CMs (AAV9-cTnT-iCre) at p0. In this system, cmKO cells are generated in neonatal CMs and identified by RFP expression. We titrated AAV vector particles and injected subcutaneously a dose of 2e10 genome copies per mouse that results in a mosaic heart with 5–10% RFP+ myocytes. The resulting RFP+ myocytes showed efficient deletion of PGC1*α*/*β*, quantified by qPCR (Figure S2A, Figure S2B, and Figure S2C). We made transverse sections and analyzed RFP+ cells with *α*-actinin staining. RFP+ cells appeared smaller than neighboring myocytes in size (Figure 3B). To precisely quantify size, we dissociated CMs using Langendorff perfusion and measured the areas of RFP+ and RFP-CMs after plating at low density (Figure 3B). RFP-cells showed a heterogeneous but progressive increase in size over time, consistent with single-cell LP-FACS cell volume analysis (Figure 3C, cyan columns). While RFP expression itself did not affect cell size (Figure S2D), we observed that RFP+ cells remain persistently smaller compared to RFP-cells. (Figure 3C, red columns). This suggests the requirement of PGC1 in cellular hypertrophy. Next, we tested whether PGC1 deficiency affects intact CM contractile function by video microscopy. RFP+ cells showed significantly lower fractional shortening and contraction velocity (Figure 3D, Figure 3E, Figure 3F, Figure S2E, Figure S2F, Figure S2G, and Figure S2H). Their contractile properties were further assessed by measuring calcium transients with the ratiometric dye Fura-2 AM. Consistent with the sarcomere shortening data, RFP+ cells showed significantly lower peak Ca2+ amplitude and slower velocity than RFP-cells (Figure 3G, Figure 3H, Figure 3I, Figure S2I, Figure S2J, Figure S2K, and Figure S2L). This kinetics suggest that PGC1-deficient myocytes develop less mature calcium cycling apparatus for Ca2+ release and re-sequestration. Together, our mosaic gene deletion approach reveals a cell-autonomous, required role of PGC1 in cellular hypertrophy and contractility development of postnatal CMs at the single cell level.

**Fig. 3.**
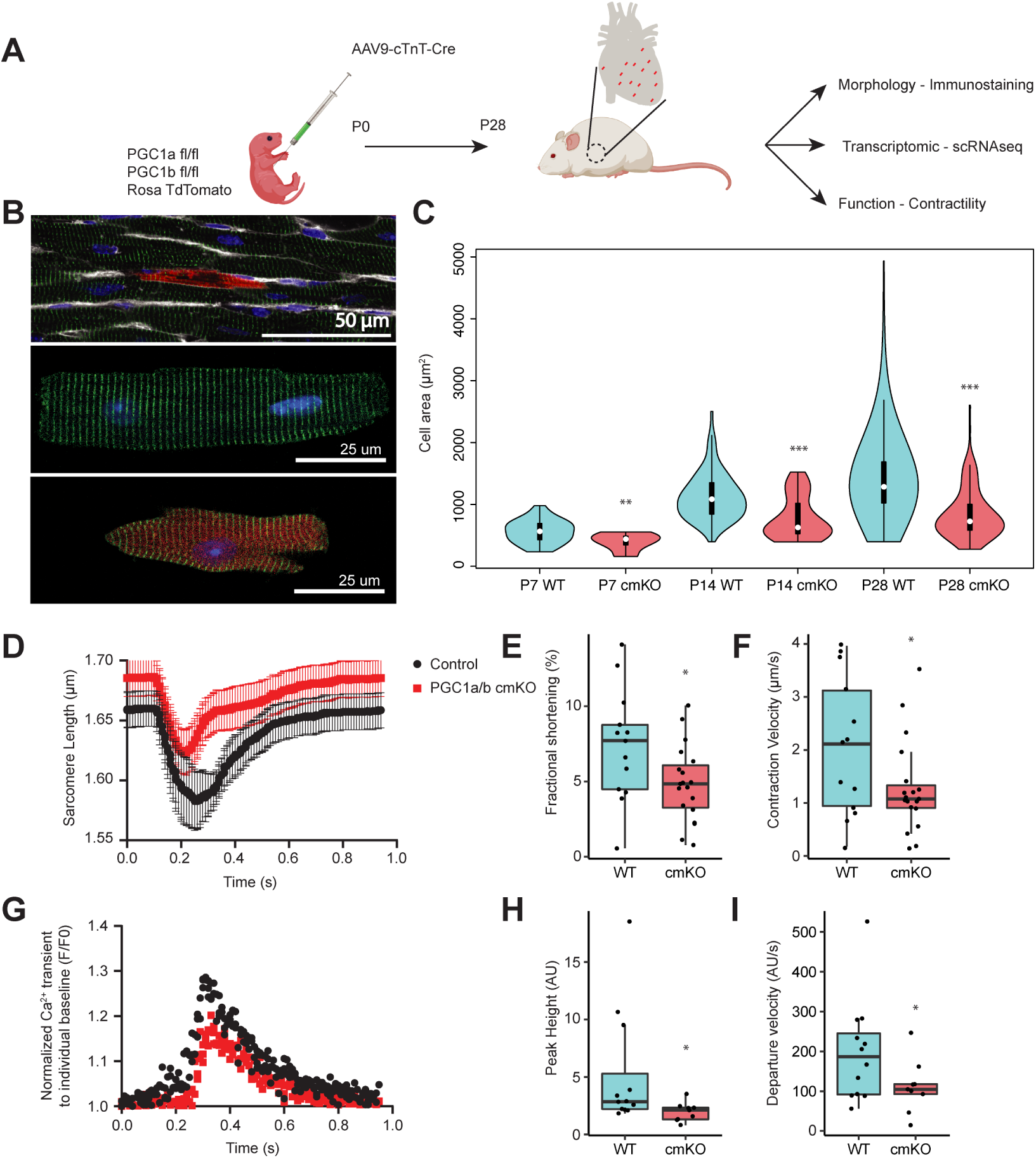
PGC1 is required for CM hypertrophy and contractility development. A, Experimental scheme showing generation and analyses of a cmKO heart achieved by injection of AAV9-cTnT-Cre into PGC1*α*/*β* flox/flox; Ai9 mice at p0. B, Heart slice showing a cmKO myocyte in myocardium (top) and dissociated control (middle) and cmKO (bottom) myocytes. C, Violin plots of cell area distributions in control (blue) and cmKO (red) CMs at p7, p14, p28. n=44,11,90,49,522,132 (left to right). D–F, Sarcomere shortening data with the average trace, fractional shortening and contraction velocity. Control n=13, cmKO n=19. G–I, Calcium handling with average calcium trace, peak height, and departure velocity. Control n=12, cmKO n=9. p-value: *<0.5,**<0.1,***<0.01.

### PGC1/PPAR*α* Regulates Genes Affecting Cell Size, Calcium Handling, and Mitochondrial Activity

Given the crucial role of PGC1 in developing myocyte hypertrophy and contractility, we investigated how PGC1 mediates these processes at the single-cell transcriptome level. After deleting PGC1 as above, we isolated RFP+ (PGC1 cmKO) cells by LP-FACS from p7–p28 hearts and conducted scRNA-seq analysis. The resulting trajectory reconstructed by Monocle with control and cmKO cells showed that PGC1-deficient CMs become less heterogeneous at p7 (Figure 4A and Figure 4B). They also maintained lower maturation scores throughout the stages (Figure 4B), which is consistent with the failure to increase in contractility and size.

**Fig. 4.**
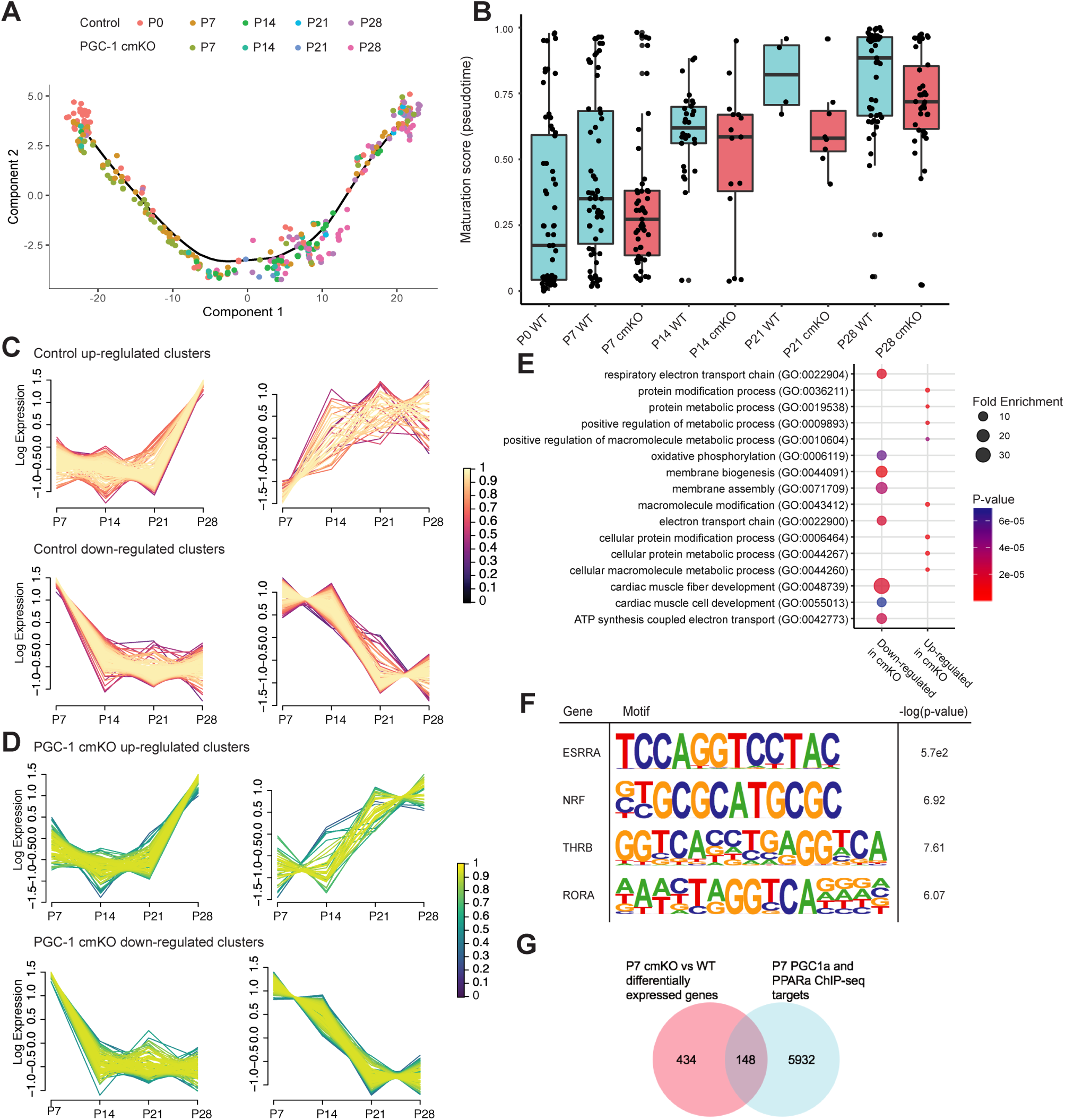
PGC1-deficiency leads to maturation defects in postnatal CMs and affects gene regulatory networks controlling muscle development and mitochondrial process. A, Single-cell transcriptomic trajectory of control and PGC1 cmKO CMs. B, Distribution of pseudotime maturation scores analyzed from p0–p28. C, Fuzzy clustering with selected clusters for upregulated (top) and downregulated (bottom) genes. Color indicates membership score of each gene in the cluster. D, Fuzzy clustering of PGC1 cmKO CMs showing upregulated (top) and downregulated (bottom) clusters. E, GO term visualization by fold enrichment (dot size) and p-value (dot color). F, Motif analysis for PGC1*α*. G, Venn diagram of differentially expressed genes in control and PGC1 cmKO CMs and PGC1*α*/PPAR*α* ChIP-seq peaks at p7.

To determine how PGC1 signaling mediates these processes, we used fuzzy clustering to group the temporal trends of individual gene expression in both control and PGC1 cmKO CMs (Figure 4C and Figure 4D). We initially selected upregulated and downregulated clusters for each group and subsequently performed overlap analysis. Notably, the analysis showed that less than 7.6% or 16.2% of genes overlap between control and PGC1 cmKO CMs in upregulated or downregulated clusters, respectively (Figure 4C and Figure 4D). This suggests that gene regulatory networks associated with maturation have been severely disrupted in PGC1-deficient CMs. GO analysis of differentially expressed genes (329 upregulated, 255 downregulated) showed that muscle fiber/cell development and mitochondrial/electron transport chain processes are significantly impaired in PGC1 cmKO cells (Figure 4E).

PGC1 binds to PPAR*α* to transcriptionally activate their target genes (Vega et al., 2000). To identify their target genes during postnatal CM development, we performed ChIP-seq on p7 hearts with antibodies targeting PGC1*α* and PPAR*α*. Antibody specificities were validated by enrichment of their known target genes (Figure S3A, Figure S3B, Figure S3C, Figure S3D, and Figure S3E). Motif analysis revealed that binding sites for ESRRA, NRF, THR*β*, and RORA are enriched in chromatin bound by PGC1/PPAR*α* which identified known and predicted binding partners (Figure 4F). We further compared genes differentially expressed in PGC1-deleted CMs with ChIP-enriched peaks in the genome and used HOMER to annotate peaks. This analysis identified 148 genes directly regulated by PGC1/PPAR*α* (Figure 4G and Table S1). They included several novel targets of PGC1/PPAR*α* with unknown roles in CM maturation, including genes encoding Yap1, a transcriptional effector of Hippo signaling with crucial roles in cardiac regeneration (Xin et al., 2013), SF3B2/SAP18, RNA splicing factors (Golas et al., 2003; Singh et al., 2010), TIMM50, a mitochondrial translocase regulating mitochondrial function (Tort et al., 2019), and STRIP1, a core component of the striatin-interacting phosphatases and kinase complex regulating cell contractility (Suryavanshi et al., 2018).

### PGC1/PPAR*α* Agonists Increase Size, Contractility, and Mitochondrial Activity of PSC-CMs

Since PGC1 is required for CM hypertrophy and contractility, and its levels and activity remain low in PSC-CMs (Uosaki et al., 2015), we further investigated if increased levels of PGC1 can promote the maturation of PSC-CMs. For this, we initially increased PGC1 levels by expressing GFP-PGC1 (Puigserver et al., 1998) in PSC-CMs. We found that GFP+ CMs became significantly larger than GFP-CMs after transfection (Figure S4A). The hypertrophic growth was recapitulated by pharmacological stimulation with pyrroloquinoline quinone (PQQ), a widely used PGC1 activator (Chowanadisai et al., 2010) (Figure 5A, Figure 5B, Figure 5F, and Figure S4C). A similar effect was also observed when cells were treated with the PPAR*α*-specific ligand WY14643 (Krey et al., 1997). Individual treatment significantly improved contractile force, measured by traction force microscopy, which was further increased when combined (Figure 5C and Figure S4B). This phenotype was accompanied with a significant increase in mitochondrial density and activity, determined by electron microscopy and Seahorse assay (Figure S4D, Figure S4E, Figure S4F, and Figure S4G). These data suggest that PGC1/PPAR*α* have an instructive role in PSC-CM maturation.

**Fig. 5.**
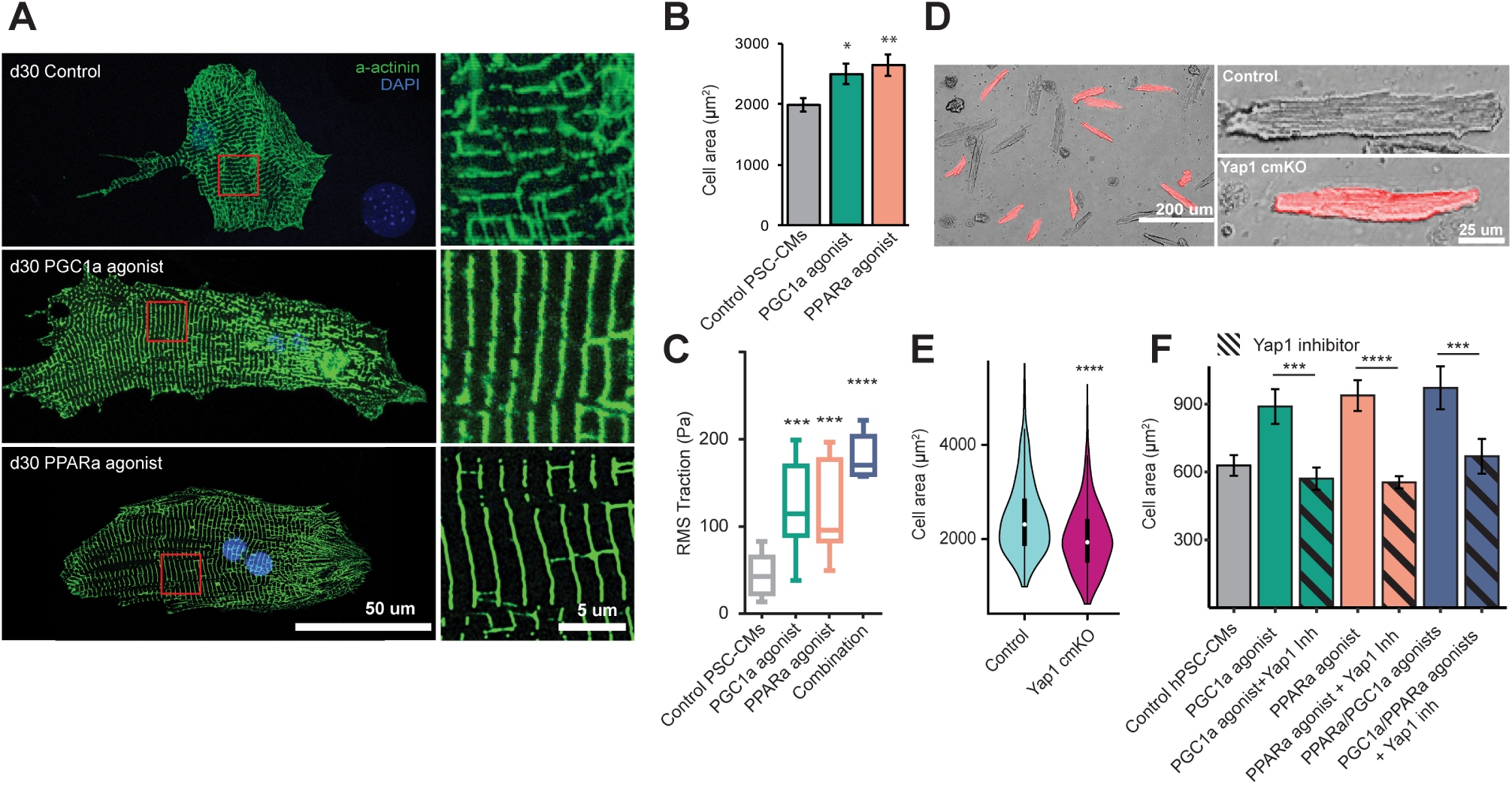
PGC1/PPAR*α* promotes PSC-CM maturation through Yap1. A, Control and PGC1*α*/PPAR*α* agonist-treated mouse ESC-CMs at day 30 (dissociated and replated at low density). The CMs were stained with *α*-actinin antibody (green) and dapi (blue). Insets show magnified views of boxed areas shown in left. PQQ 10*µ*M, WY14643 1*µ*M. B, Quantification of cell area of mouse ESC-CMs treated with PGC1*α*/PPAR*α* agonists for 15 days. Control n=192, PGC1 agonist n=162, PPAR*α* agonist n=118. C, Fourier transform traction microscopy showing RMS traction in pascals of control and treated mouse ESC-CMs. D, Representative images of control (RFP-) and Yap1 cmKO (RFP+) CMs isolated from p32 hearts. E, Quantification of cell area of control and Yap1 cmKO CMs. Control n=319, Yap1 cmKO n=404. F, Cell area measurements for human ESC-CMs treated with PGC1*α*/PPAR*α* agonists in combination with Yap inhibitor ((R)-PFI-2 1*µ*m). n=25, 26, 41, 52, 40, 31, 22 by column.

### PGC1/PPAR*α* Signaling Promotes CM Maturation by Regulating Key Upstream Regulators of Cellular Hypertrophy and Calcium Handling

To determine how PGC1/PPAR*α* mediates CM growth, we analyzed expression levels of conserved cell size regulators Mtor, Yap1, Igf1. Among these, we found that Yap1, one of the validated PGC1/PPAR*α* targets by ChIP-seq, is markedly downregulated in PGC1 cmKO cells (Table S2). To test if Yap1 affects CM size in vivo, we generated Yap1 cmKO cells in postnatal hearts as shown in Figure 2E and analyzed their growth. Notably, the cmKO CMs became significantly smaller than normal CMs (Figure 5D and Figure 5E), indicating that Yap1 may mediate PGC1/PPAR*α* signals for CM hypertrophy. To test this, we chemically blocked Yap1 transcriptional activity with the Yap1 inhibitor (R)-PFI-2 (Barsyte-Lovejoy et al., 2014) in PSC-CMs stimulated with PGC1/PPAR*α* agonists. Blocking Yap1 activity indeed abolished PGC1/PPAR*α*-mediated cell growth (Figure 5F). These data suggest that Yap1 is required for PGC1/PPAR*α* to promote CM hypertrophy.

Since PGC1/PPAR*α* signals promoted CM contractility, we sought to identify downstream genes regulating calcium handling. To do this, we performed a single-cell high-throughput functional assay with PSC-CMs (Cunningham et al., 2017; McKeithan et al., 2017; Yu et al., 2018) treated with PPAR*α*-specific ligands as shown in Figure 6A. Single cell analysis of calcium handling revealed that ligand-treated PSC-CMs have shorter (30ms) calcium transient duration (CTD) as compared to vehicle treated cells (DMSO) (Figure 6B and Figure 6C).

**Fig. 6.**
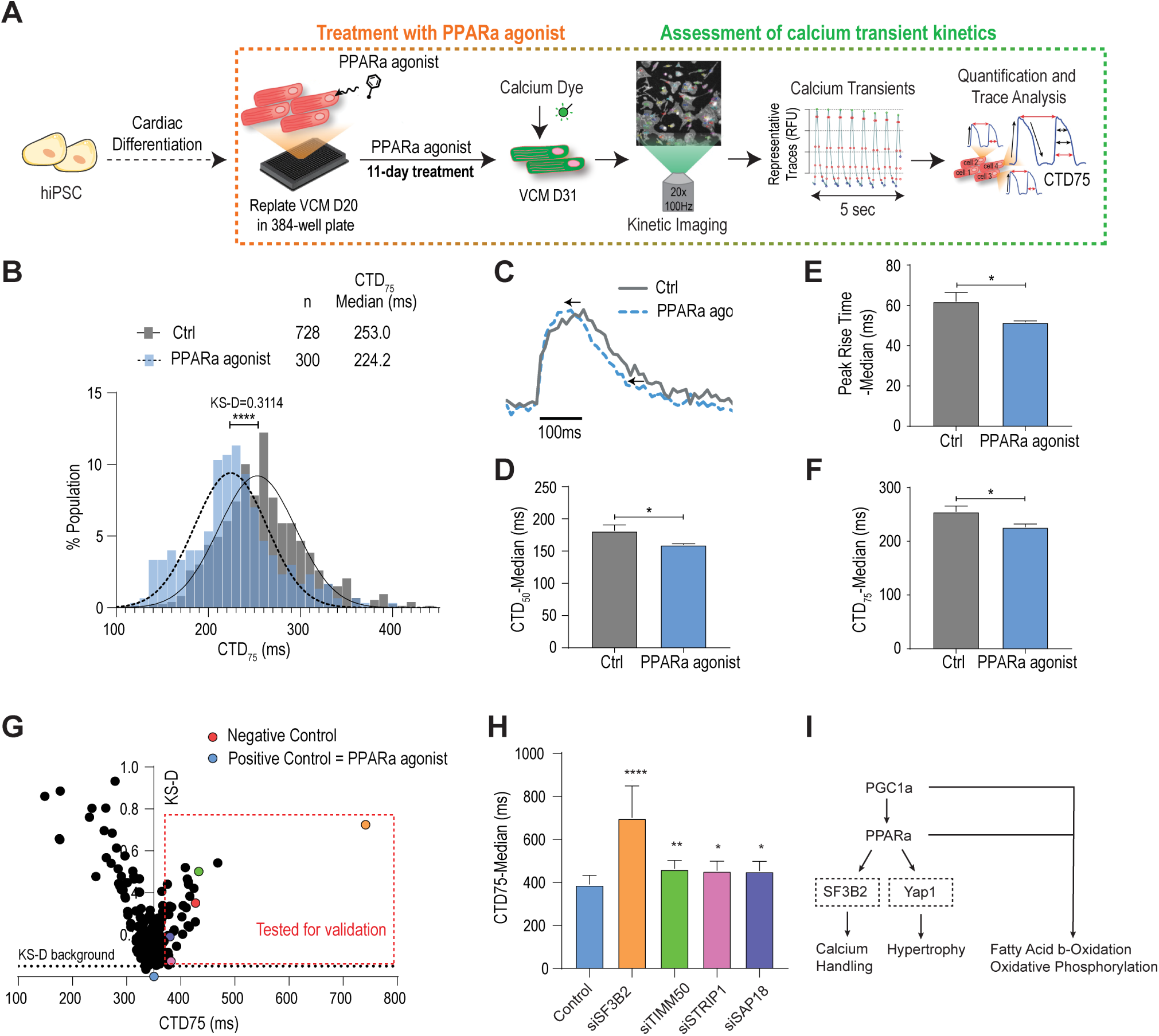
PGC1/PPAR regulate calcium handling via SF3B2. A, Experimental diagram of human ESC-CM differentiation, agonist treatment, and calcium function analyses. B–F, Distribution of calcium transient duration (CTD) 75 with sample trace and median peak rise and CTD50 and CTD75 times. G, SiRNA screen results with kernelized stein discrepancy (KS-D) and CTD75. Untreated or PPAR*α* agonist-treated ESC-CMs were used as negative or positive control, respectively. H, Median CTD75 for validated siRNAs. I, Working model. p-values: *<0.5,**<0.1,***<0.01,****<0.001. ANOVA using student’s t-test with Bonferroni correction.

Notably, calcium transient peak rise time was shorter, and CTD50 and 75 were decreased (Figure 6D, Figure 6E and Figure 6F), thereby suggesting that calcium handling properties are enhanced in ligand-treated cells. Next, to identify downstream effectors mediating the CTD shortening, we stimulated PSC-CMs with PPAR*α* ligands and applied a library of siRNAs (4 siRNAs/gene) targeting 148 genes directly regulated by PGC1/PPAR*α* signaling (Figure S5A). Our results showed that the ability of PPAR*α* signaling to shorten CTD is significantly impaired when targeting SF3B2/SAP18, TIMM50, and STRIP1 (Figure 6G, Figure 6H, Figure S5B, Figure S5C, Figure S5D, Figure S5E, Figure S5F, and Figure S5G). This finding suggests that these genes mediate PGC1/PPAR*α* signaling for the improvement of calcium handling in PSC-CMs.

## Discussion

In the present study, we investigated how individual CMs give rise to mature cells, a fundamental, yet poorly understood event. We have demonstrated that CMs are heterogeneous in transcriptome and size during postnatal stages, when significant levels of adaptive changes occur, with variable expression levels of genes responsible for cell growth and contractility. We found that PGC1 signaling, whose activity is increased until adulthood in vivo but remained low in PSC-CMs, is necessary to regulate genes crucial for hypertrophy and contractility in addition to mitochondrial genes, albeit in a heterogeneous manner among cell populations. These findings provide fundamental and mechanistic insights into the postnatal maturation of single CMs, coordinated by PGC1 signaling (Figure 6I).

Earlier transcriptome analysis of developing hearts suggested that CMs develop unidirectionally towards a more mature state through discrete developmental stages (Uosaki et al., 2015), yet it remains unknown if this represents homogeneous maturation of individual myocytes. It is intriguing that CMs exhibit and maintain a high level of developmental heterogeneity throughout postnatal maturation stages. Indeed, a small subset of early myocytes showed mature transcriptomes as early as P7, whereas some late myocytes still have transcriptomes resembling immature cells. This suggests that postnatal CMs mature at different rates, and achieving full maturation of individual myocytes may not precisely follow the developmental timeline or may not occur in all myocytes. This is supported by heterogeneous cell size and the presence of myocytes expressing Myh7 and Tnni1 at P28, considered not to be expressed in mature myocytes. It would be important to further investigate where the immature or mature cells are located and if the immature cells represent a small subset of proliferative myocytes present in adult hearts.

Understanding the factors and mechanisms underlying cardiac maturation is of great importance, but there is very little information available at this point. This could be attributed to the difficulty of staging and defining milestones of the process that takes place over a long period of time. In fact, a recent study demonstrated the importance of stage-specific gene regulation in CM maturation (Guo et al., 2018). Our study suggests that PGC1 is upregulated after birth and its activity is required and sufficient to promote CM maturation. Intriguingly, while PGC1 signaling is known to play a critical role in controlling mitochondrial biogenesis and cellular metabolism (Finck and Kelly, 2006), our single-cell analysis showed that it regulates multiple aspects of cellular events, including cell size, calcium handling, and contractility in addition to oxidative phosphorylation, during cardiac maturation. This is particularly surprising given its conserved and universal role in energy metabolism. These findings suggest that PGC1 signaling may function as a master regulator in postnatal CM maturation.

A number of previously unrecognized genes were found targeted by PGC1/PPAR*α* in postnatal stages. In particular, splicing factors SF3B2/SAP18 were required for functional maturation of PSC-CMs. RNA splicing is an important post-transcriptional mechanism, and its abnormal regulation is closely associated with human diseases including heart disease (Faustino and Cooper, 2003; Mirtschink et al., 2015). While splicing factors were shown to promote neuronal maturation (Jacko et al., 2018), their role in the context of CM maturation remains to be determined. We also found that PGC1/PPAR directly regulates Yap1, a conserved transcriptional regulator of organ/cell size (Lloyd, 2013). The role of Yap1 has been extensively studied on CM proliferation for heart repair (Wang et al., 2018), but growing evidence suggests its critical role in cellular hypertrophy (Perez-Gonzalez et al., 2019; Windmueller and Morrisey, 2015). Similarly, our data showed that Yap1 is a key mediator of PGC1/PPAR for CM growth. A previous study suggested that Yap1 is not required for physiological cardiac hypertrophy (von Gise et al., 2012), but this discrepancy could be due to methodological or technical differences in cell size quantification. Further investigation would be necessary to precisely dissect the effect of Yap1 on CM hypertrophy.

PSC-CMs have great potential for a wide range of preclinical and clinical applications, including cardiac disease modeling, drug discovery, and regenerative medicine. The resulting myocytes, however, are prematurely arrested at late embryonic stage in culture (DeLaughter et al., 2016; Uosaki et al., 2015), and the inability to produce mature myocytes from PSCs is a major hurdle for their broad applicability. We found that PGC1/PPAR signaling remains inactive even in long-term-cultured PSC-CMs, and its activation promotes their size, contractility, and metabolism, key features of myocyte maturation. This suggests that the developmental arrest is at least in part attributed to the lack of PGC1/PPAR activity in PSC-CMs. This is consistent with the structural and functional arrest observed in PGC1 KO CMs in vivo. Knowing the complexity of cardiac maturation, our finding is expected to help us further investigate gene regulatory networks and barriers controlling the distinct processes of cardiac maturation.

## Methods

### Animals and PSC Culture

PGC1*α*/*β* flox, Ai9, Yap1flox mice (Lai et al., 2008; Lin et al., 2004; Madisen et al., 2010; Zhang et al., 2010) were obtained from the Jackson Laboratory. All protocols involving animals followed U.S NIH guidelines and were approved by the animal and care use committee of the Johns Hopkins Medical Institutions. CMs were differentiated from mouse and human ESCs (E14 and H9, WiCell) as described (Cho et al., 2017b). Briefly, mouse ESCs were cultured in gelatin-coated flasks with stem cell maintenance media (GMEM + 10% FBS with 3 um Chir99021, 1 um PD98059, glutamax, non-essential amino acids, and sodium pyruvate). To differentiate, embryoid bodies were formed by plating 80,000 cells/mL into uncoated dishes. The medium used for differentiation and CM culture contained 75% IMDM, 25% Ham’s F12 (Cellgro) with B27 without vitamin A, N2, BSA, glutamax, Penicillin/Streptomycin, L ascorbic acid, and a-monothioglycerol. 48h later, embryoid bodies were collected and induced for 48h with Bmp4 and Activin A, then dissociated and replated with XAV939 (Sigma).

### Heart dissociation, sorting, and immunostaining

Harvested hearts were placed in a Langendorff setup and perfused with a Type II Collagenase and Protease digestion buffer and stepped up with Calcium to 1 mM. Viable CMs were sorted using LP-FACS (Kannan et al., 2019). The Union Biometrica COPAS Flow Platform was used to select CMs based on extinction, time of flight, and red fluorescence. For staining, hearts were flash frozen into O.C.T Compound blocks (Fisher Healthacre 23-730-571) and stored at −80 C. Samples were sliced into 10 um sections using a cryo-microtome and placed on glass slides. Primary antibodies against *α*-actinin (Abcam) and PGC1*α* (Abcam) were used along with DAPI and Wheat Germ Agglutinin counterstains.

### Calcium handling and shortening analysis

Sorted cells were plated onto laminin-coated glass slides then loaded with Fura2AM Ca2+ dye. They were analyzed with the IonOptix imaging system and IonWizard software as described (Cho et al., 2017a).

### scRNA-seq and ChIP-seq analysis

For scRNA-seq, individual cells were fluorescently sorted into 96 well plates using the Union Biometrica COPAS Flow Platform, which were frozen on dry ice and stored at −80 C. We used SCRB-seq to prepare libraries as done (Kannan et al., 2019; Soumillon et al., 2014). Paired-end reads were sequenced using an Illumina NextSeq500 then mapped with STAR and featureCounts-based zUMIs (Parekh et al., 2018). Monocle 2 and Seurat v2.4 were used for analysis. For ChIP-seq, hearts were isolated, washed and minced then fixed in 1% paraformaldehyde crosslinking buffer. Tissue chunks were homogenized, and DNA was fragmented using 20 cycles of 30 second on then 30 seconds off of 50% power sonication. Protein G magnetic beads were incubated with the sample for 1h. Antibody (PGC1*α* Santa Cruz, PPAR*α* Abcam ChIP-grade ab227074) was incubated at 4C overnight. Protein G magnetic beads were incubated for 1h then pulled down. Chromatin was eluted from the beads then uncrosslinked by incubating overnight at 65°C. DNA was purified using phenol-chloroform following RNase and Proteinase K treatment. Libraries were prepared using Clontech DNA SMART kit (Takara Bio 634866) and sequenced on a HiSeq 4000. Reads were trimmed using trimread. We used Macs2 (Zhang et al., 2008) to call peaks and HOMER to annotate peaks. Peaks were visualized using bedtools and integrated genome viewer.

### Mitochondrial functional assay

Respiration rates were measured with Seahorse XFe96 Analyzer. CMs were plated at 1.5×104 cells per well of a 96 well XF96 Cell Culture Microplate (Aligent Technologies) and cultured for 3 days. One hour before the assay, medium was changed to RPMI without phenol red supplemented with sodium pyruvate. The Seahorse Extracellular Flux Assay Kit was used with the Mito Stress Test protocol. Inhibitor final concentrations were Oligomyocin (2.5*µ*M), FCCP (1*µ*M), and Rotenone (2.5*µ*M) + Antimycin A (2.5*µ*M).

### Electron Microscopy

D30 PSC-CMs were fixed by 2% GA in 0.075 cacodylate with 5mM MgCl2 overnight at 4°C. Samples were rinsed 3 times for 15 minutes with a 3% sucrose buffer then treated with 2% osmium and 1.5% KFeCN6 for 2 hours at 4°C. They were rinsed with 0.1 M maleate buffer pH 6.2 with 3% sucrose 3 times for 10 minutes then 2%UA in maleate/sucrose buffer for 1 hour without light. Samples were dehydrated in an ethanol ladder stepping up from 30% to 100% at 5 minutes each. They were treated with propylene oxide then EPON resin with catalyst overnight with rocking then EPON resin with catalyst for 2 hours then placed in a 60C oven for 48 hours. Sections were cut using a Diatome diamond knight collected on 2×1mm formvar-coated slot grids and stained with uranyl acetate followed by lead citrate. A Hitachi H-7600 TEM operating at 80 kV. An AMT XR-50 CCD was used to digitize images.

## Acknowledgements

We thank Deepthi Ashok and Dr. Brian O’Rourke for helping mitochondria assays, Danielle Rigau for animal work assistance, George McNamara for confocal imaging assistance, Michael Delannoy for electron microscopy assistance, and Dr. Daniel Kelly for providing technical assistance for ChIP. We thank Dr. Leslie Tung for helpful discussions and critical feedback. This work was supported by grants from NIH, MSCRF, AHA, and JHU TMTM.

## Author Contributions

S.M., M.M. designed and carried out this work. S.K. helped with LP-FACS and scRNA-seq analysis. A.K, A.C. designed and performed high-throughput PSC-CM assays. B.L., D.K., D.A.K. provided expertise in contractility assays. E.T. designed and performed hPSC-CM analysis. S.P., S.A. performed traction microscopy. S.K. performed animal work and image analysis. P.A. helped with designing in vitro screen. R.Z. helped with scRNA-seq data analysis. H.U. provided conceptual input. C.K. designed and supervised this work and wrote the manuscript with S.M.

---

**Fig. S1.**
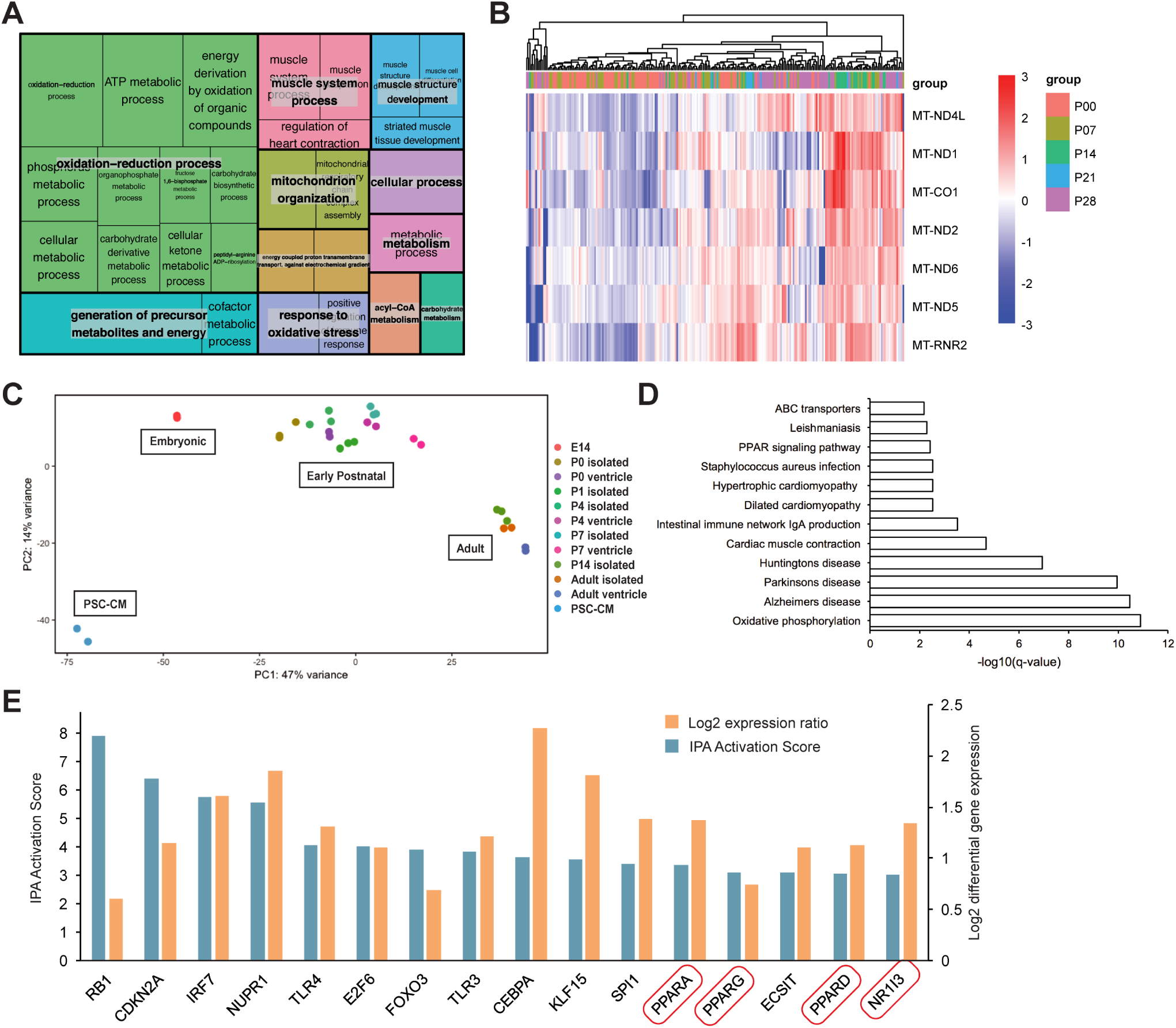
scRNA-seq clustering and meta-analysis of bulk RNA-seq. A, Treemap of GO terms of differentially expressed genes from P0 to P28 with box size representing the −log10 p-value. B, Heatmap of mitochondrial gene expression with hierarchical clustering of time-points. C, PCA plot of two componentions of bulk RNA-seq datasets of cardiac maturation (GSE64403, GSE47948, GSE95762, GSE79883). D, KEGG pathway mapping of differentially expressed genes from neonatal to adult CMs. E, Top IPA transcriptional regulators with log fold change and IPA activation score visualized. PPAR family nuclear receptors are among the top hits.

**Fig. S2.**
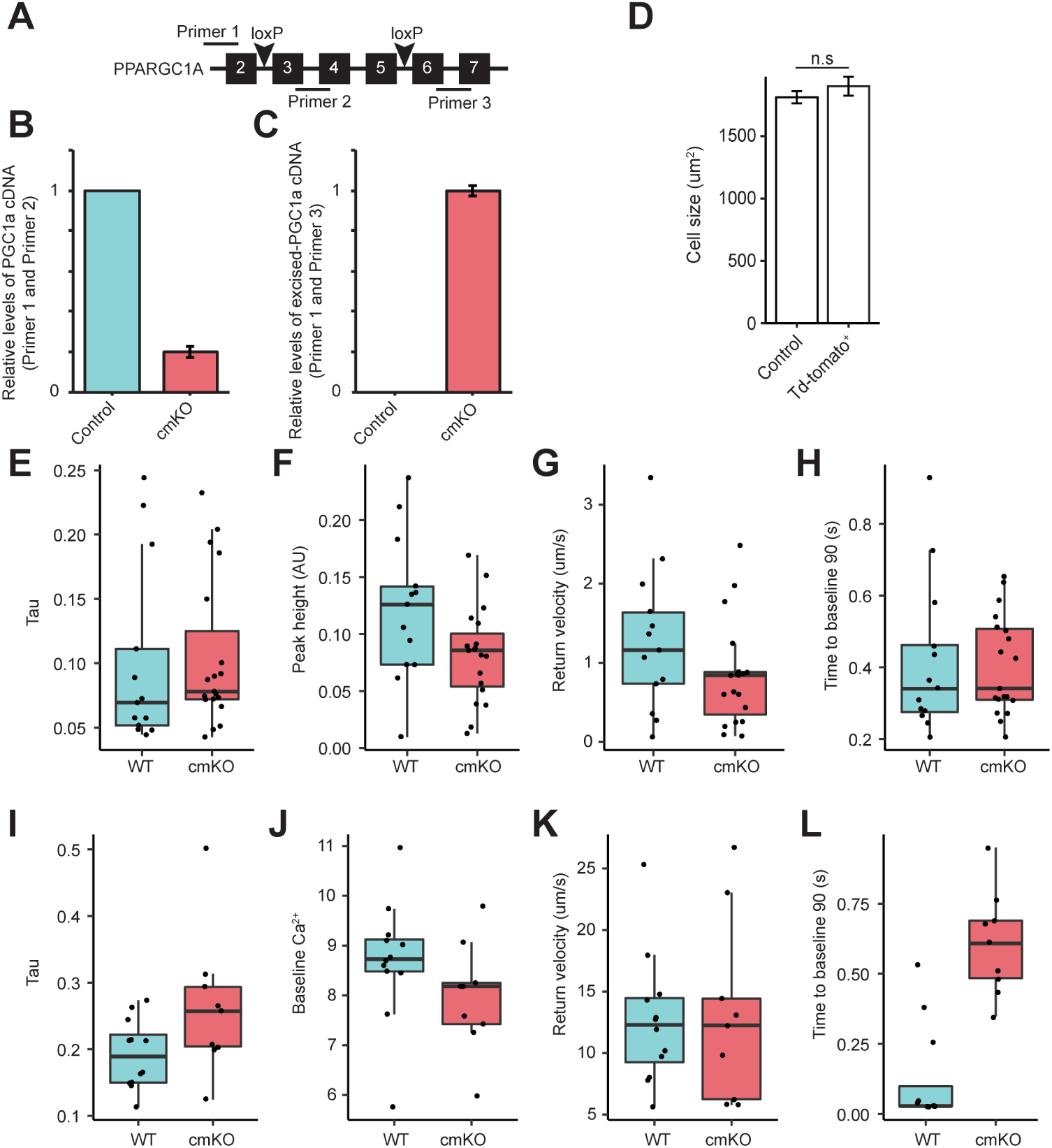
PGC1 cmKO validation and additional functional metrics A, Diagram showing exons 2–7 of PGC1*α* and the use of qPCR primers to quantify levels of PGC1 in control and cmKO CMs. B, Normalized levels of PGC1*α* in control and cmKO CMs. C, Normalized levels of excised PGC1*α*. D, Cell size measurements for myocytes isolated from p28 hearts of Ai9 mice injected with AAV9-cTnT-Cre at p0. E–H, Contractility parameters for control and PGC1 cmKO CMs. control n=13, cmKO n=19. I–L, Calcium transient parameters for control and PGC1 cmKO CMs. control n=12, cmKO n= 9.

**Fig. S3.**
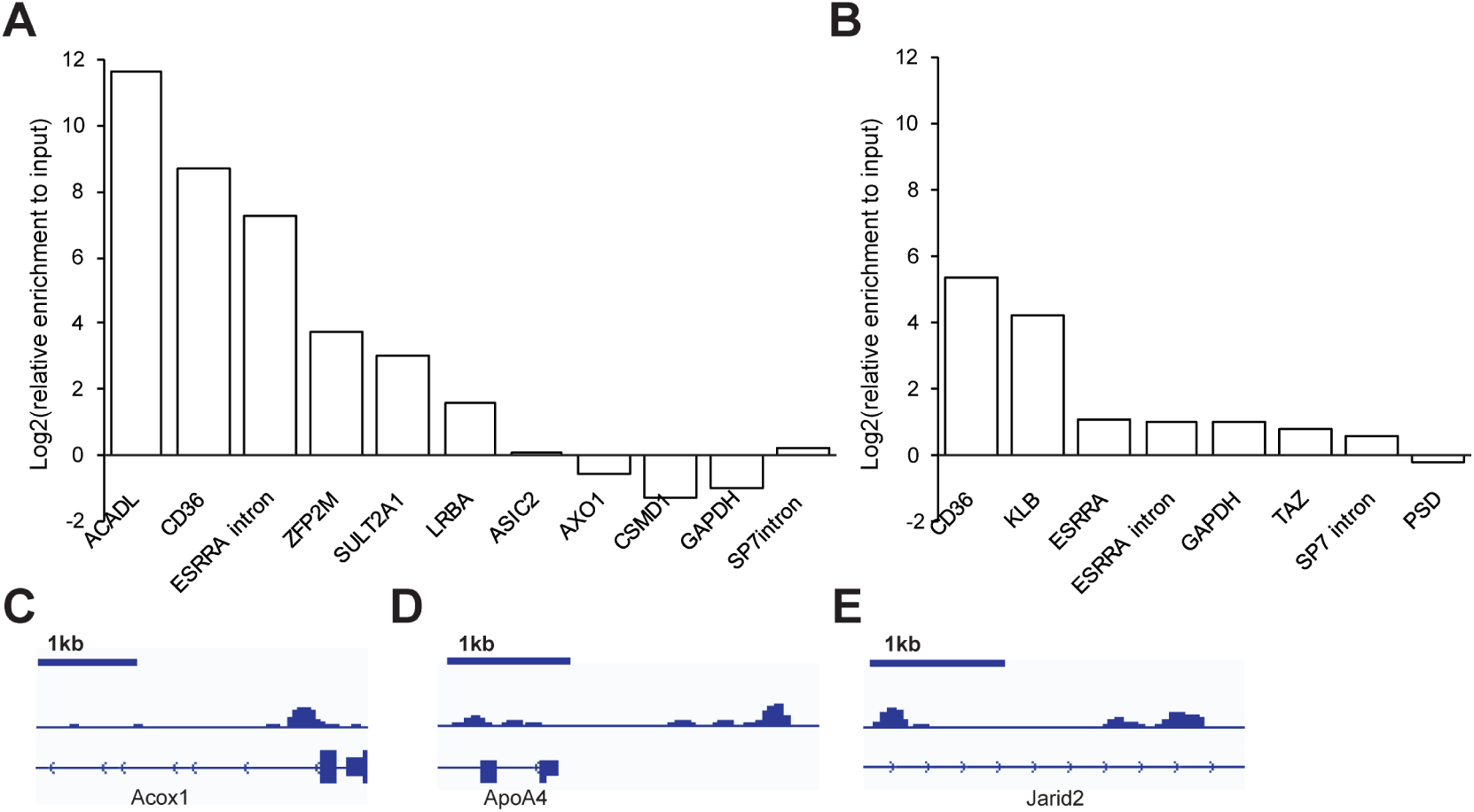
ChIP-seq antibody validation. A, B, ChIP-qPCR showing log fold relative enrichment of promoter sequences of PPAR*α* (A)/PGC1 (B) targets and control DNA regions (GAPDH, SP7 intron, ESRRA intron). C, D, Peaks of known binding sites for PGC1*α* (C) and PPAR*α* (D). e, peak showing novel binding site for PGC1 in intronic region of Jarid2.

**Fig. S4.**
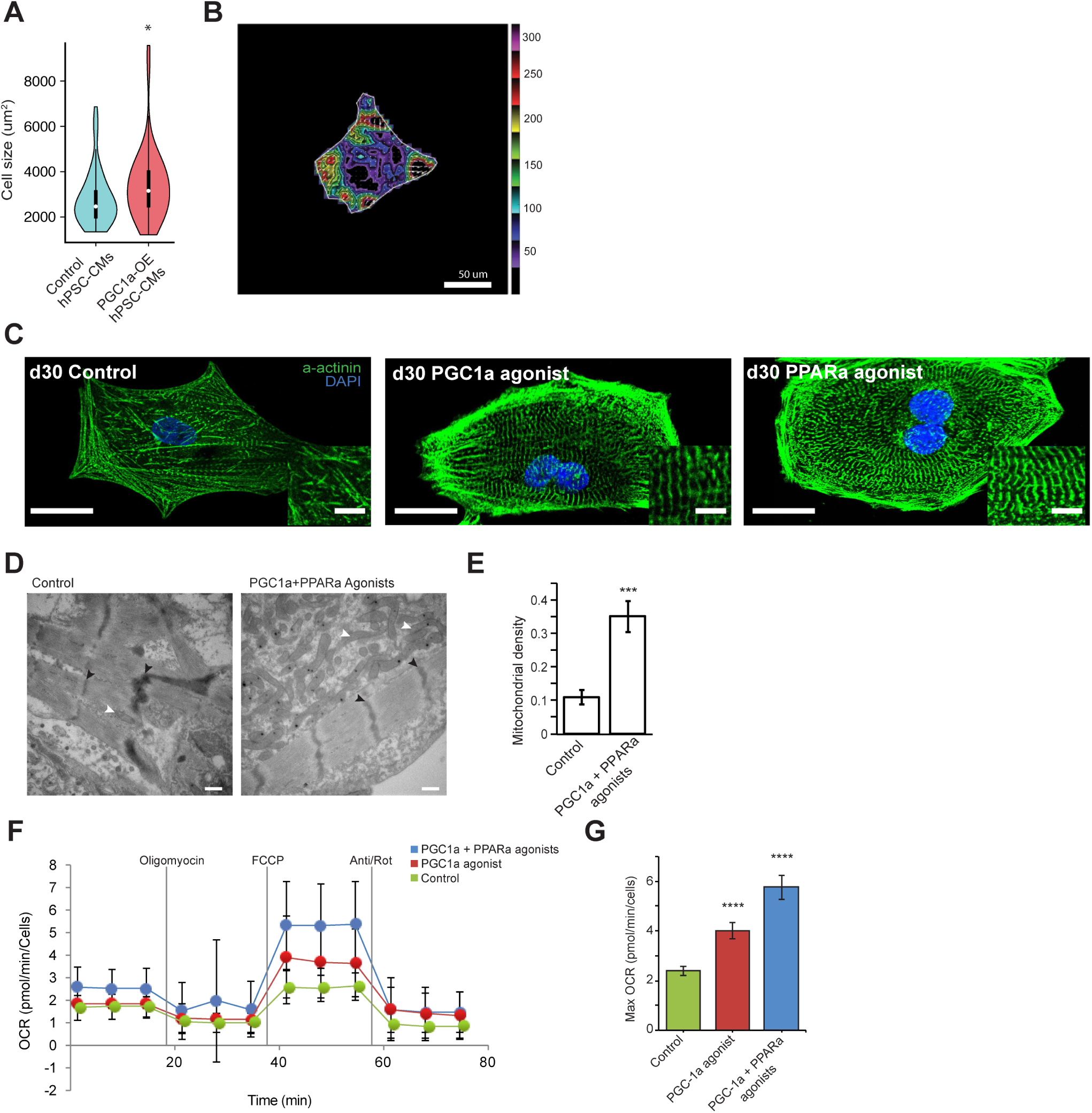
PGC1/PPAR activation increases cell size, contraction, and mitochondrial activity. A, Cell area quantification of human ESC-CMs transfected with control and GFP-GFP-PGC1 expression construct, measured 4 days after transfection. control n=55, GFP-PGC1 n=46. B, Fourier transform traction force microscopy showing force vectors during ESC-CM contraction. C, Images of day 30 human ESC-CMs treated with PGC1/PPAR*α* agonists for 2 weeks after differentiation. D, Electron microscopy images of control and PGC1/PPAR*α* agonist-treated day 30 ESC-CMs. Scale bar = 500 nm. White arrowheads show mitochondria and black arrowheads show z-bands. E, Mitochondrial density quantified from EM images. F, Seahorse XF96 measurements of oxygen consumption rate in control and agonist-treated d30 ESC-CMs.

**Fig. S5.**
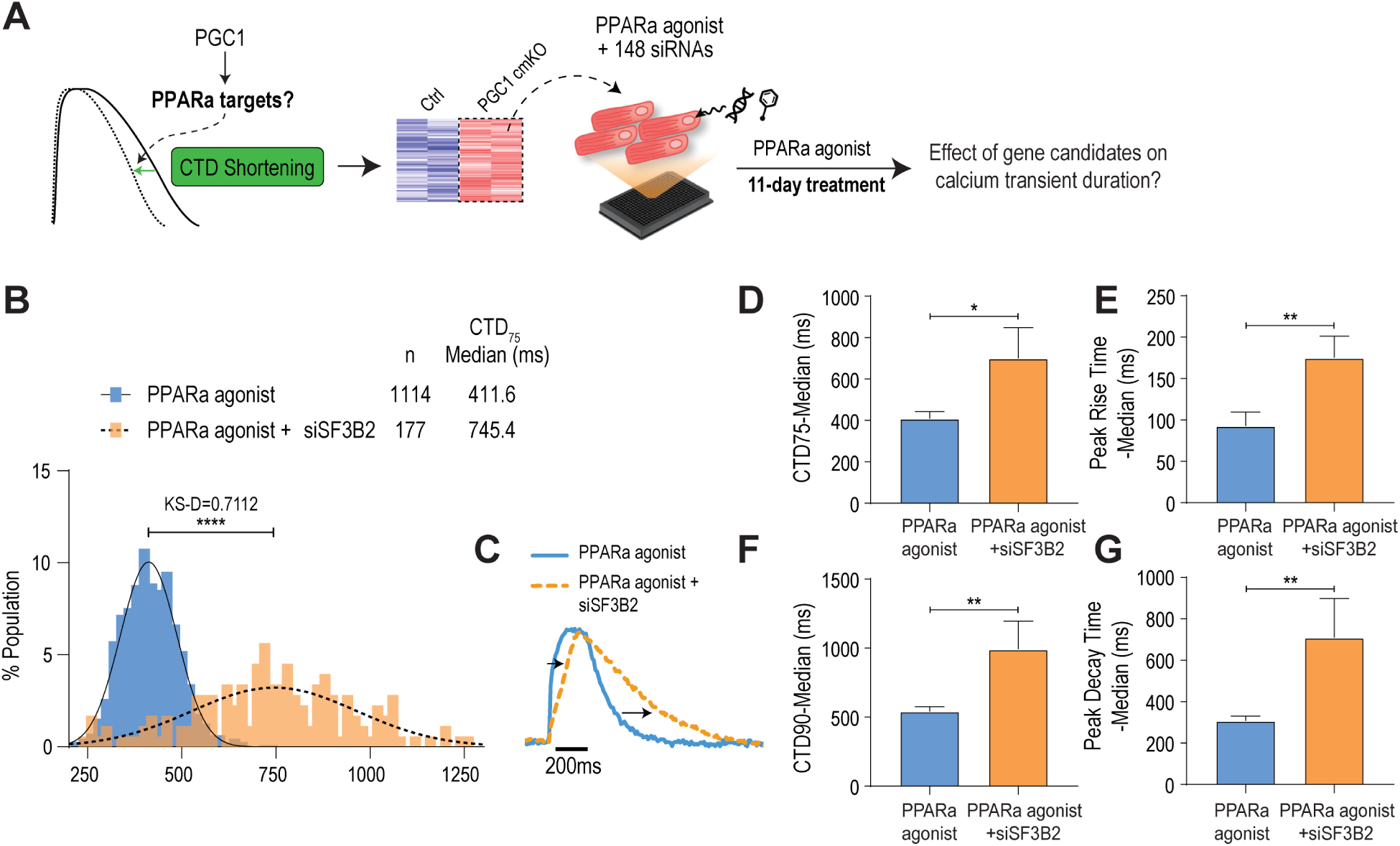
siRNA-based assay for PGC1/PPAR targets driving calcium handling maturation. A, Experimental design for siRNA-based functional assay. B, Distribution of CTD75 in PPAR*α* agonist-treated human ESC-CMs with/without siRNA targeting SF3B2. C, Representative traces of ESC-CM calcium transient. D–G, Calcium transient parameters showing functional effects of SF3B2 knockdown.

